# Glocal dendritic spikes: variation in branch activity during dendritic spiking indicates multiple functional units in the pyramidal neuron apical tuft

**DOI:** 10.1101/2022.03.15.484426

**Authors:** Erez Geron

## Abstract

There is a lively debate on how many functional units exist within the dendritic tree of pyramidal neurons (PNs). The classical view that the PN’s dendritic tree comprises several computational units ^1-4^ was challenged by recent in-vivo studies that identified mainly widespread dendritic spikes ^5-10^, suggesting a low degree of dendritic compartmentalization. We reasoned that global spikes do not rule out dendritic compartmentalization if these spikes contain a high variation in branch activity. Therefore, we turned to image Ca^2+^ activity in many apical dendrites from individual layer (L) 5 PNs of the primary motor cortex (M1). For proper dendritic morphology and Ca^2+^ activity analysis, we used mice with sparse double labeling of PNs with a structural marker (tdTomato) and a GCaMP6s probe. We first collected data for complete registration of the apical tuft dendritic structure. We then used 3D two-photon microscopy to image Ca^2+^ activity within these tufts during treadmill running. Using these, we imaged up to 30 apical branches from individual L5 PNs, including branches separated by as many as 11 bifurcations and more than 1000 µm of intracellular distance. We found that local spikes were rare, but global spikes often displayed heterogeneous Ca^2+^ elevation across branches, giving rise to an activity pattern across the tuft. This spiking pattern was dynamic and according to unsupervised hierarchical cluster analysis, fitted to 2-6 distinct patterns, even over short imaging periods. Taken together, we find a considerable variation in branch activity within and between global dendritic spikes, supporting multiple dendritic functional units.

**Graphical abstract:** 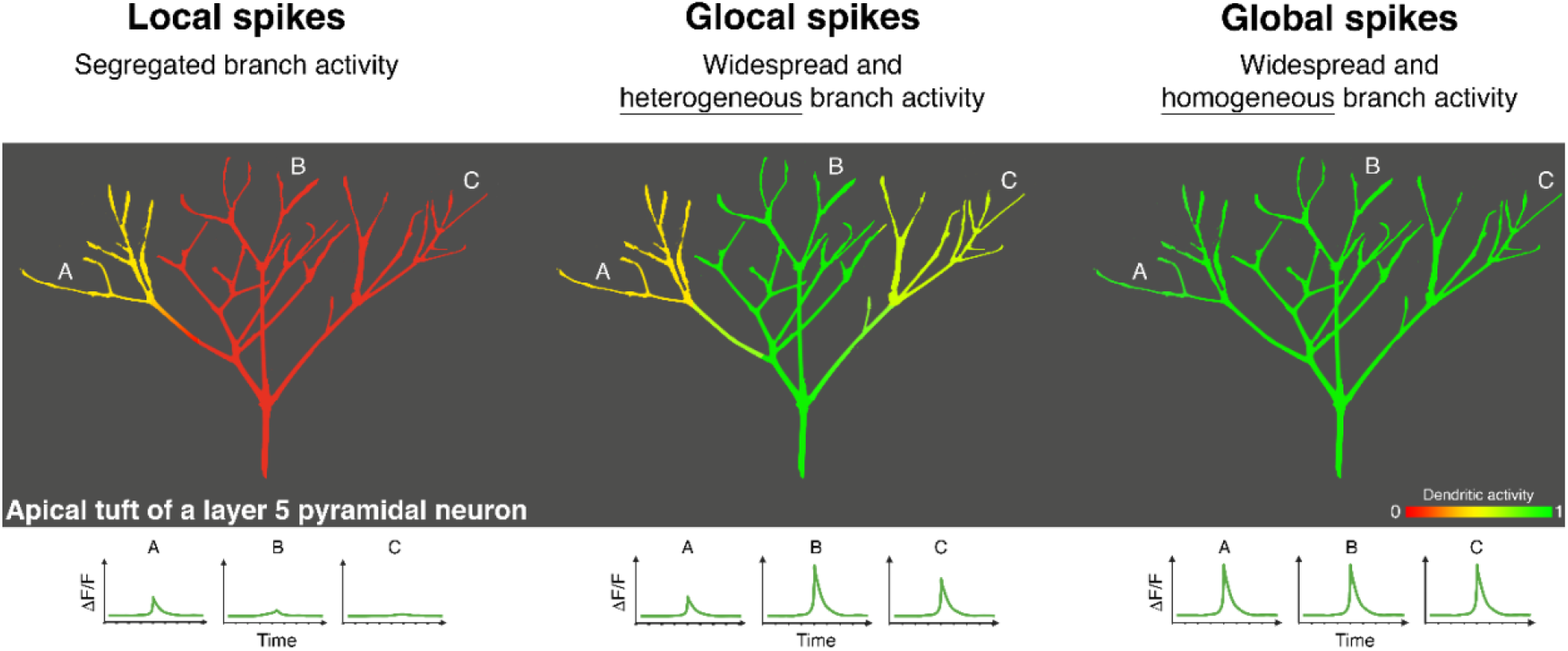

We detected three main spike types. Local spikes were evident in a minority of the branches and had low amplitude. Contrastingly, global spikes often displayed homogeneous levels of branch activity and were the strongest and most prolonged spikes. An intermediate pattern was the “glocal spike,” where a global spike exhibited heterogeneous branch activity. Spikes with heterogeneous Ca^2+^ elevation across branches constituted at least a quarter of all spikes. In the social sciences, the term “glocalization” refers to global phenomena that are adjusted locally, such as food items sold worldwide but attuned to local tastes ^11^. Influenced by that, we refer to global spikes with heterogeneous branch activity as “glocal spikes.”

## Introduction

A large portion of the excitatory input of pyramidal neurons (PNs) is received onto apical dendrites located hundreds of microns away from the soma. Despite this distance, distal inputs can profoundly influence somatic output through dendritic Ca^2+^ spikes ^12^. Dendritic Ca^2+^ spikes are non-linear electric events that involve a large and sustained membrane depolarization and cytosolic Ca^2+^ elevation, which can be triggered separately of somatic activity ^12-14^. Dendritic integration through spikes allows neurons to act as two-layer processors and enhance their computational power. Furthermore, a large amount of computational and *in-vitro* studies suggested that groups of dendrites within individual cells can act separately of one another, and thus render neurons as multi-layer processing units ^1-4^. Although viewing neurons as multi-layer units gained vast support across decades, it has been challenged by recent studies that assessed dendritic compartmentalization in vivo.

Dendritic compartmentalization is often inferred by the prevalence of spikes, i.e., the fraction of active branches within a tuft during a spike. The abundance of widespread (global) spikes indicates low activity variability and suggests that the apical tuft operates as a single-layer unit. In contrast, a predominance of compartmentalized dendritic activity (local spikes) supports several processing units. Notably, there is no clear definition of local or glocal spikes. For example, local spikes have been defined as those detected in a single branch ^15^, a few branches ^16^, and as spikes detected in the tuft but not in the trunk/soma ^10^. Inference of compartmentalization is further complicated by studying a few branches of individual cells and using GECIs with limited sensitivity, both likely to underestimate dendritic compartmentalization ^17,18^. Albeit, most recent *in vivo* studies have reported a predominance of global activity ^5-9,19-21^ (but see ^16,22^). Similarly, most studies that monitored dendritic and somatic activities simultaneously have reported high dendritic-somatic coupling, suggesting low dendritic compartmentalization ^7,8,10,20^ (but see ^23^). These studies have led to the proposal that dendritic activity is less compartmentalized than previously thought and that apical tufts operate predominantly as a single computational unit ^24^.

We studied the variation in dendritic Ca^2+^ spikes by imaging apical dendrites from individual L5 PNs using *in-vivo* volumetric two-photon microscopy. L5 PNs in the primary motor cortex (M1) were sparsely labeled with a structural marker and a GCaMP probe, which enabled analysis of dendritic Ca^2+^ activity in tufts whose morphology was well defined. Importantly, our volumetric imaging allowed for scanning of a continuous volume (40 µm), which enabled monitoring the activity of up to tens of dendrites from an individual PN. During training with a treadmill running task, both local and global dendritic Ca^2+^ spikes were detected. Although local spikes constituted a small portion of all spikes, spikes with partial prevalence were common: half of the spikes were detected in 75 % or less of the branches. Notably, Ca^2+^ elevation during single global spikes differed between branches, leading to a specific activity pattern across the tuft. This dendritic activity pattern was dynamic and changed between spikes. Unsupervised cluster analysis identified 2-6 distinct activity patterns across spikes. Therefore, widespread and variable dendritic spikes are consistent with the existence of a number of computational units within the PN apical tuft dendrite.

## Results

### An AAV infection strategy for dendritic imaging of individual layer 5 pyramidal neurons

Monitoring of Ca^2+^ activity from sub-cellular structures, such as dendrites and spines, benefits from high signal to noise ratio, which is facilitated by sparse neuronal labeling. Sparse labeling was achieved by gating GECI expression by a low titer of a *Cre*-harboring virus that activated the expression of in a small percentage of the infected population. We calibrated our infection setup to allow for labeling of neuronal structure in addition to monitoring Ca^2+^ activity. Here, mice were injected with a mixture of three AAVs, where a low titer of a Cre-harboring virus (AAV.*CaMKII*::Cre; final titer of 10^8^ VG/mL (diluted 1:100,000; see Materials and Methods)) enabled the expression of Cre-dependent tdTomato and GCaMP6s. This approach resulted in sparse double-labeling, enabling functional imaging in branches whose relation to other branches was known (**Figures 1A-C**). Importantly, virus injection was performed in neonates ^25^ (**Figure S1**), which circumvented any detectable changes of skull and meninges when evaluated four weeks after infection (**Figure 1A**). Furthermore, neonatal injection did not result in inflammation within infected and surrounding cortical regions as microglia displayed a normal tile appearance in Cx3cr1::eGFP mice injected during the same neonatal period (**Figure S2**). Together, these data indicate that this infection strategy is highly beneficial for repeated *in-vivo* dendritic imaging.

**Figure 1.**
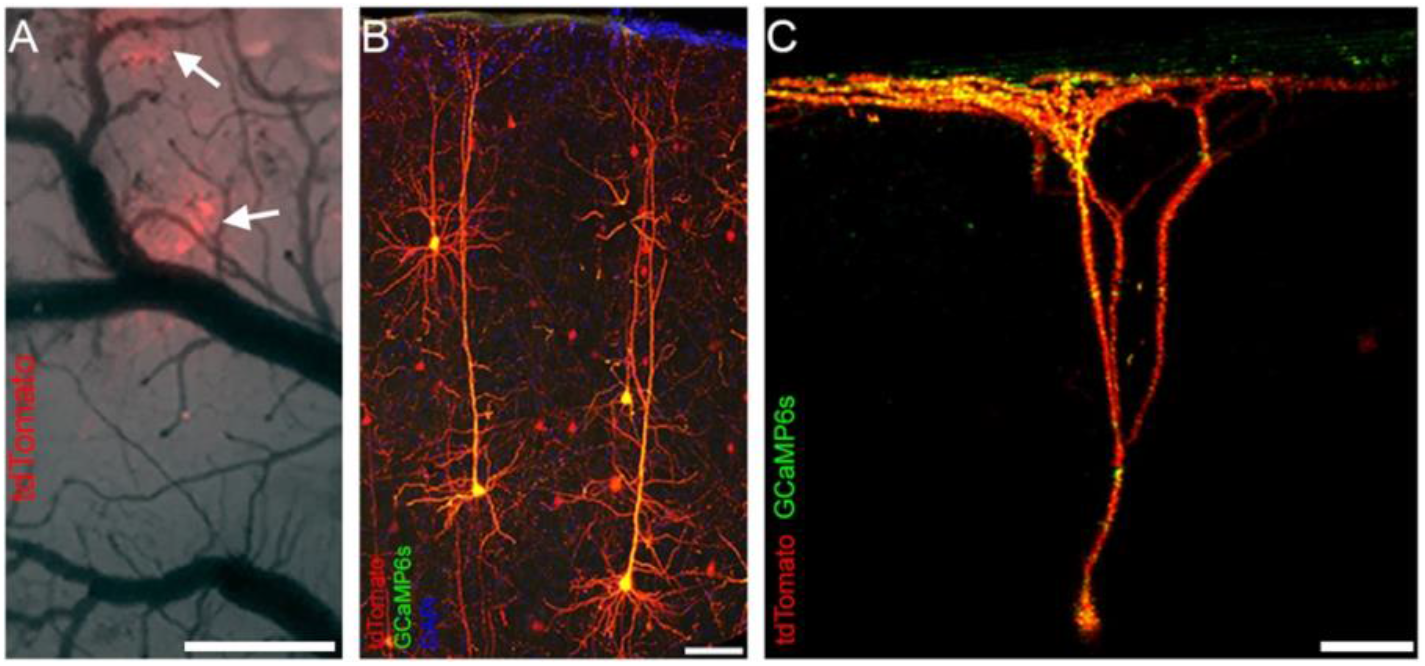
Labeling individual layer 5 pyramidal neurons by AAV infection of neonate mice. A-C. Examples of sparse double-labeling of pyramidal neurons (PNs) with tdTomato (red) and GCaMP6s (green). Mice were injected with AAVs harboring Cre, tdTomato, and GCaMP6s constructs at P1-3 to label PNs, and imaged at P30. (A) Top view of a cortical area of ∼10 mm^2^ bearing tufts of two labeled PNs (arrows). (B) Side view of a cortical slice with several double-labeled PNs. The slice was stained with DAPI after fixation. (C) 3D rendering of multiple z stacks acquired in an awake animal. Scale bars represent 1 mm in a, and 100 μm in b and c.

### Spike prevalence during training: local spikes were rare but partial prevalence was common

In the experiments presented in figures 2-4, we first collected high-resolution z-images of the dendritic arbor of individual L5 PNs in awake mice, using conventional two-photon microscopy. We then trained these mice with a treadmill running task while imaging the dendritic Ca^2+^ activity with volumetric microscopy (**Figures 2A, B**; **Videos 1-3**). Importantly, correlating structure and activity using sparse double labeling allowed for measuring dendritic activity within tufts whose morphology was well defined. Our volumetric setup allowed for scanning of a continuous volume of 40 μm (x × y × z: 1000 × 1000 × 40 μm; 80 z planes, at a speed of 2 Hz; located 20-60 μm below the pial surface) ^26^. With this method, we were able to obtain data from a large number of dendrites per neuron (up to 27 dendritic branches from a single cell, 14.4 ± 2.04, Ave ± SEM; 130 branches, 9 neurons from 8 animals) (**Figures 2A**; **S2**; **Videos 1-3**; **Table S1**). These included branches that were relatively far away from one another: the most-distant branch pairs were separated by an average of 9.1 ± 0.33 bifurcations and by an intracellular distance of 633.1 ± 72.1 μm (mean ± SEM; distance measured between bifurcations; 9 neurons. Note, that this distance can be greater than the distance of these branches from the soma) (**Table S1**). The bifurcation order of imaged branches ranged from 2^nd^ to 7^th^. Most of the branches were 4^th^ or 5^th^ order (34.3 ± 4.1 and 38.1 ± 2.6 %, respectively). Finally, scanning a continuous volume that is ten times larger than that of conventional two-photon microscopy reduces the relative contamination induced by movements in the z-axis We first characterized the frequency of local and global spikes. Importantly, the definitions of local and global spikes vary between studies and research groups. For example, local spikes have been defined as those detected in a single branch ^15^, a few branches ^16^, and as spikes detected in the tuft but not in the trunk/soma ^10^. Here, we monitored dendritic activity only within the tuft; therefore, as a working definition, we refer to spikes detected in 25 % or less of the branches as local (prevalence score ≤ 25 %) and to spikes detected in more than 75 % of the branches as global. When we applied these criteria, local spikes constituted only 12.5 % of all spikes (40/318) (**Figure 2C**). A similar value was reached when prevalence was calculated for each neuron (15.1 ± 4.04; mean ± SEM; 9 neurons) (**Figure 2D**). Global spikes constituted 54 % of all spikes (172/318), and 51.6 ± 4.04 % when calculated for each neuron (mean ± SEM) (**Figures 2C, D**). Although local spikes were rare, partial prevalence was common: nearly half of the spikes were not global (46 %; 146/318) (**Figures 2C, D**; **Table S1**). As expected, stronger spikes were detected in more branches, and these two variables correlated positively (r= 0.52; P< 0.0001, n= 318 spikes, 9 neurons) (**Figure 2E**). Taken together, a considerable portion of the spikes displayed partial prevalence and did not fall under the typical classification of local or global spikes.

**Figure 2.**
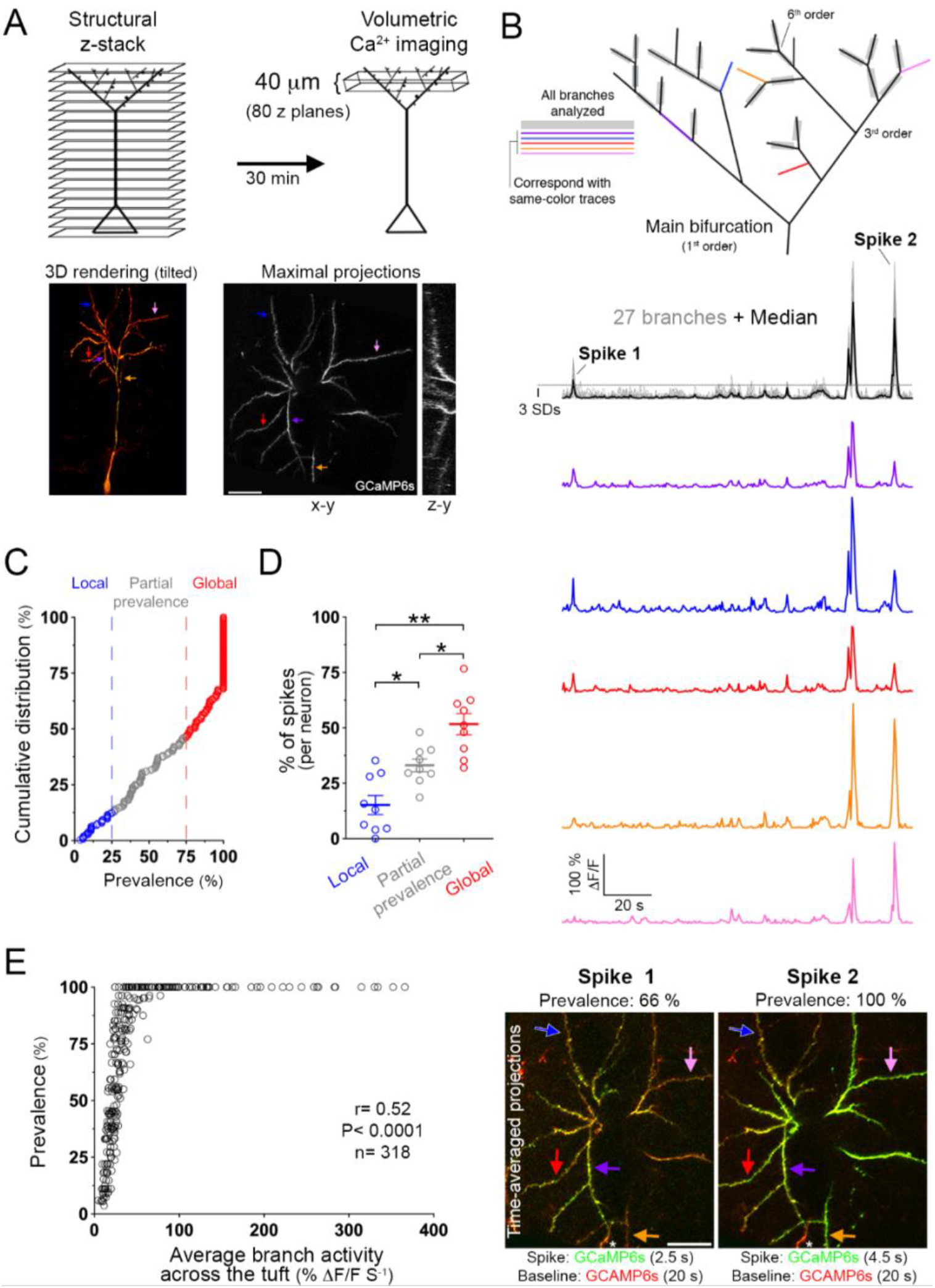
Spike prevalence during training: local spikes were rare but partial prevalence was common. A. We first collected high-resolution z-stacks of single L5 PNs in M1 of awake mice to later reconstruct their morphology. The animals were then trained with a forced motor running task while dendritic Ca^2+^ activity was imaged using volumetric two-photon microscopy. Note that arrows of the same color in the 3D rendering point to corresponding branches in the maximal projections in A, and in the arbor schematics and ΔF/F traces in panel B. Asterisks mark a dendrite from a neighboring cell. Scale bar in the lower-right panel represents 40 μm. B. Images and ΔF/F traces of dendritic spikes with different prevalence. *Top*: Schematics of the branching order of the apical dendrites of the PN in panel A. ΔF/F traces of Ca^2+^ activity of the five colored branches are presented below. Branches marked in gray were analyzed. Center: ΔF/F traces of Ca^2+^ activity of the indicated branches (color-coded in top panel). *Bottom*: Time-averaged projections of two spikes. GCaMP6s fluorescence was averaged across the spikes’ durations (green; 2.5 and 4.5 s for spikes 1 and 2, respectively) and overlaid on a projection of the baseline (20 s, red, same for both images). Color intensities in both images are to scale. Scale bar in the lower-left panel represents 40 μm. This panel corresponds with Video S1. C, D. (C) Prevalence (% of active branches) of all 318 spikes detected in this study (each circle represents a spike). Thick and thin black lines denote the median and the quartiles, respectively. Dashed red and blue lines denote the criteria used for global and local spikes, respectively. (D) Prevalence calculated for each neuron (n= 9; from 8 animals). (Each circle represents the portion of the classified spikes in a single cell). *< 0.024, **= 0.0029 (one-way ANOVA; Fisher’s LSD test). E. Prevalence as a function of spike strength (r= 0.52; P: < 0.0001; n=318 spikes; 9 neurons from 8 animals).

### Glocal spikes: global spikes with heterogeneous branch activity

We noted that differed in their Ca^2+^ elevation during a single spike during a spike (**Figure 3A**; **Video 2**). Such variation in branch activity was most apparent during local spikes but was common also in global ones. We derived the spikes’ coefficient of variation (CV) to characterize the variability in branch activity during single spikes. The coefficient of variation is a normalized measure of variability that indicates the relative size of the standard deviation from the mean (SD/mean). Figure 3A depicts traces from 18 branches and focuses on two spikes with CV scores of 87.3 and 41.03 % (images on the left are time-averaged projections of the spikes’ entire duration). This study’s average CV score of all spikes reached 46.5 ± 1.1 % (mean ± SEM; 318 spikes from 9 neurons). A similar score was achieved when calculated across neurons (48.8 ± 3.3 %; mean ± SEM) (**Table S1**). Notably, when the CV was plotted as a function of prevalence, a strong negative correlation was evident (r= −0.57, P< 0.0001) (**Figure 3B**). Furthermore, within global spikes, the spikes’ CV correlated negatively with their amplitude (r= −0.28, P= 0.0001, 172 spikes) (**Figure 3C**).

**Figure 3.**
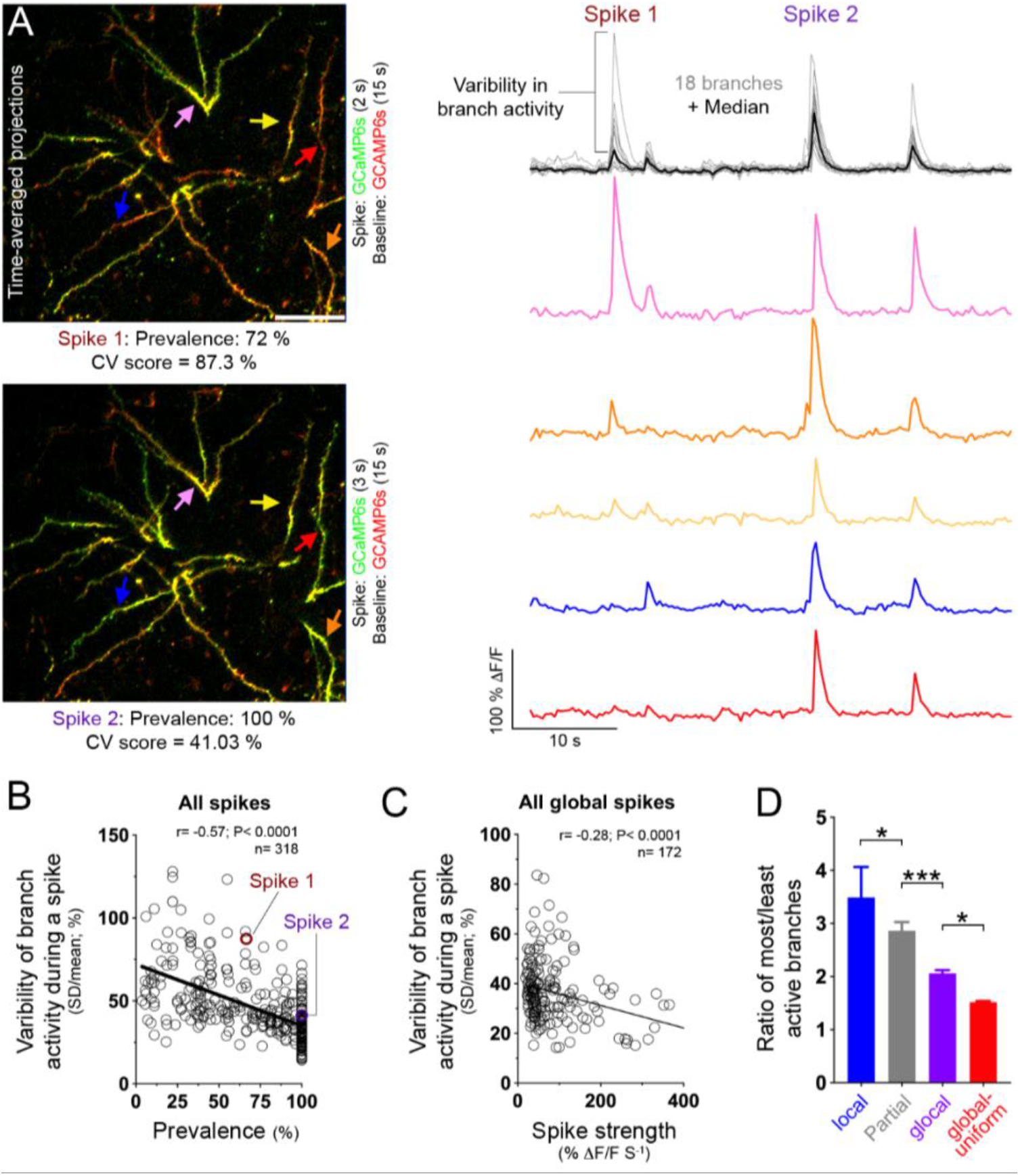
Glocal dendritic spikes: global spikes with heterogeneous branch activity. A. Images and ΔF/F traces of dendritic Ca^2+^ activity across a tuft. Time-averaged projections of two spikes (left). GCaMP6s fluorescence was averaged across the entire duration of the spikes (green; 2 and 3 s for spikes 1 and 2, respectively) and overlaid on a projection of the baseline (15 s, red, equal for both spikes). The color intensity in both images is to scale. Colored arrows correspond with same-color traces on the right. Scale bar in the upper panel represents 50 μm. This panel corresponds with Video S2. B, C. (B) Variability of branch activity during a spike as a function of the spike’s prevalence (r= −0.57; P< 0.0001; 318 spikes from 9 neurons in 8 animals). Variability of branch activity was measured by the spike’s coefficient of variation (a normalized measure of variability that calculates the relative size of the standard deviation from the mean (SD/mean)). (C) Variability of branch activity during a spike as a function of the spike’s strength, for all global spikes (r= −0.28; P< 0.0001; 172 spikes; 9 neurons). D. Activity ratio of the most and least active branches (top and bottom 25%). Spikes were classified according to their prevalence and their branch variability to: local (prevalence ≥ 25 %), partial prevalence (prevalence 25 % < > 75 %), glocal (prevalence ≥ 75 %; CV ≥ 35 %), and global-uniform (prevalence ≥ 75 %; CV< 35 %). *< 0.04, ***= 0.0008 (one-way ANOVA; Fisher’s LSD test).

In social sciences, the term glocalization refers to widespread phenomena that are adjusted locally (e.g., food items sold worldwide but attuned to local tastes) ^11^. We adopted this term to describe global spikes with high variability in branch activity. Here, glocal spikes were defined as those with above-average branch variability, and global-uniform spikes as those with below-average variability (average variability: CV= 36.5 %, 172 spikes).

To grasp better the variability in branch activity during single spikes, we also calculated the ratio between the activity of the most active (top 25%) and least active (bottom 25%) branches. Notably, this ratio differed significantly between all spike groups (local, partial prevalence, glocal, and global-uniform) (**Figure 3D**). Collectively, we detected a considerable variation in branch activity during single spikes. This variability was evident not only in local spikes but also in spikes with global and partial prevalence. This mixture of broad prevalence and local differences in branch activity prompted us to refer to such spikes as glocal.

### Distinct activity patterns across spikes

By itself, variation in branch activity during spiking is not indicative of multiple dendritic units. If spikes consistently display the same branch activity pattern, the tuft could be considered a single-layer unit, where some branches contribute more than others to the dendritic output. However, suppose the activity pattern changes between spikes. In that case, distinct branches would dominate different spikes, strongly indicating several dendritic processing units. With this in mind, we characterized the variation in activity patterns between spikes. For example, Figure 4A depicts time-averaged images and ΔF/F traces of 16 branches from an individual L5 PN, focusing on three spikes with 100 % prevalence. For simplicity, we first display the activities of four branches (marked with colored arrows) across the three spikes. It is evident that a very active branch in a particular spike is not as active in other spikes (orange trace in spike 2 vs. spikes 1 and 3) (**Figure 4A**, right; **Video 3)**. Indeed, correlating activity across all branches during the three spikes revealed that spikes 1 and 3 were more similar to the one another than to spike 2 (Pearson’s coefficient: r_1↔3_= 0.75; r_1↔ 2_ = 0.101; r_2↔ 3_= 0.33) (**Figure 4A**; **Video 3**).

**Figure 4.**
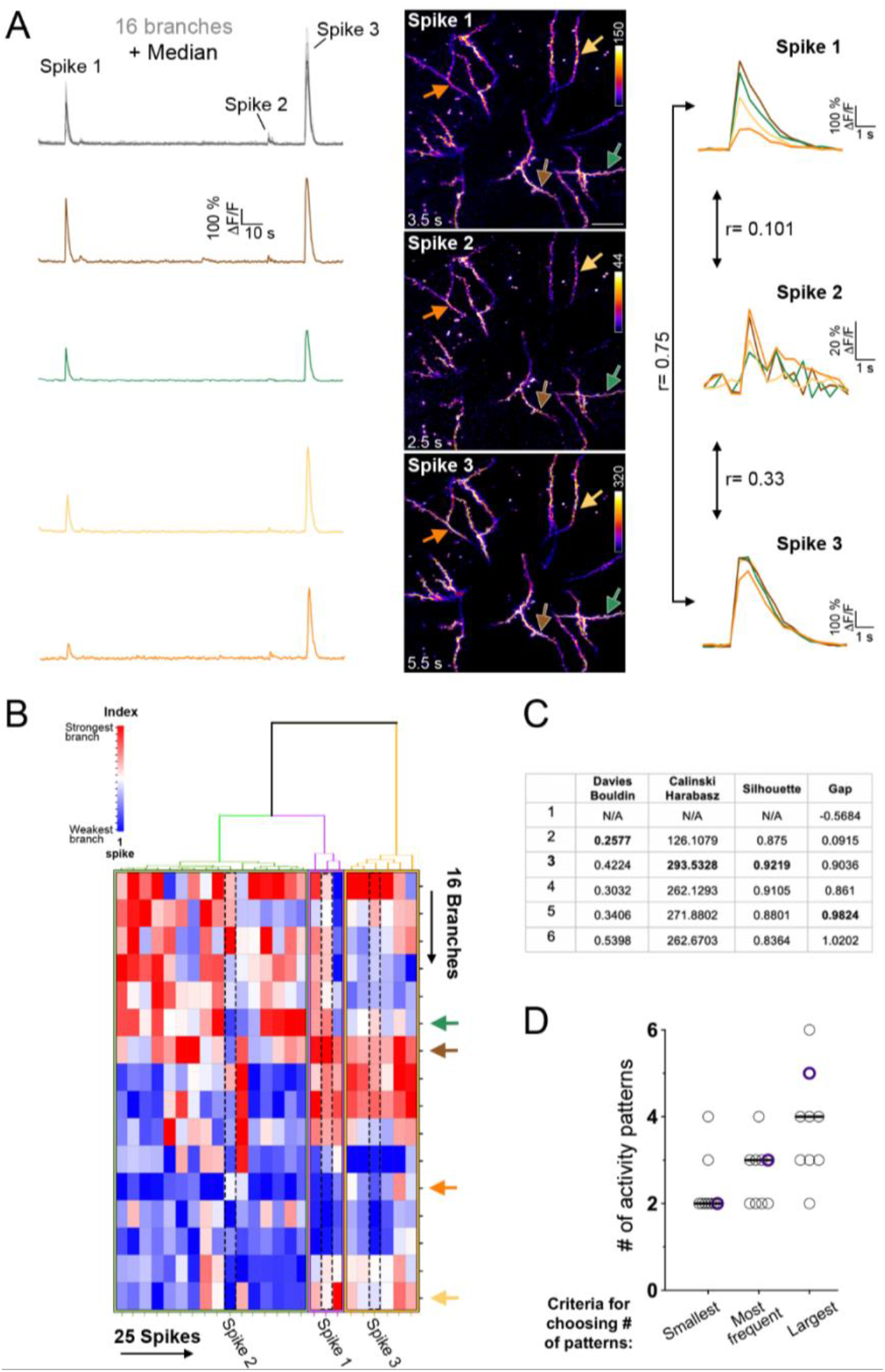
Dynamic patterns of branch activity between spikes. **A.** Images and ΔF/F traces of dendritic Ca^2+^ activity across a tuft. *Left*: ΔF/F traces of Ca^2+^ activity of 16 branches from an individual L5 PN tuft. The colored traces are of the branches marked with same-color arrows in the center panel. *Center*: Time-averaged projections of three spikes. GCaMP6s fluorescence was averaged across the entire duration of the spikes. Numbers above the intensity bars denote the maximal branch activity (% ΔF/F). Numbers at the lower left corners denote the spikes’ duration. Scale bar in the upper panel represents 40 μm. *Right*: ΔF/F traces of the four indicated branches during spikes 1-3. Correlation between the spikes was calculated for 16 branches. This panel corresponds with Video S3. **B.** Unsupervised cluster analysis of the spikes of the neuron presented in A (25 spikes (columns); 16 branches (rows)). Spikes were clustered according to branch activity (Ward’s method). Colored arrows, and numbered spikes, correspond with those in A. **C.** Estimation of the optimal number of activity clusters by four clustering evaluation indices. Estimation is for the spikes presented in B. Bolded values indicate the optimal number of clusters according to each of the indices. **D.** Number of distinct activity patterns according to different criteria. Purple circles denote the number of activity patterns for the neuron displayed in panels A-C (n=9 neurons).

We next wanted to assess how many distinct activity patterns can be differentiated within the spikes of a given neuron (35.3 ± 6.5 spikes per neuron; imaged over 19.8 ± 1.3 min; mean ± SEM, 9 neurons). To this end, we performed an unsupervised cluster analysis in which spikes were clustered according to their branch activity (Ward’s method) (**Figure 4B**). In general, when an exploratory cluster analysis is performed, a key issue is whether the identified clusters reflect the true functional organization of the acquired data. To address this, we used four cluster evaluation indices to estimate the optimal number of clusters ^27^. Importantly, one of these indices (the gap index) is suited for detection of a single cluster ^28^. As a conservative estimate, we first display the smallest number of clusters identified across the four indices (**Figures 4C, D**). This yielded an estimate of 2-4 distinct activity patterns (median= 2; 9 neurons). When the largest number of clusters was considered, 2-6 distinct activity patterns emerged (median= 4) (**Figures 4C, D**). These analyses support the idea that PN apical dendrites possess the modularity to enable distinct spiking patterns across different branches.

## Discussion

We studied the variation in dendritic Ca^2+^ spikes in apical tufts of L5 PNs of the motor cortex during training with a treadmill running task. Using sparse double labeling and volumetric imaging of a continuous volume, we were able to analyze the activity of a large number of dendrites from individual L5 PNs. We found that apical tufts displayed high variability in many spiking parameters: strength, duration, prevalence, and pattern. Ca^2+^ elevation across the tuft was not uniform but varied between branches during single spikes. In addition, variation in branch activity was not restricted to local spikes but was also common in global spikes. This mixture of global prevalence and locally increased Ca^2+^ elevation prompted us to use the term “glocal spikes.” Finally, the pattern of branch activity across the tuft was dynamic between spikes. Unsupervised cluster analysis identified 2-6 activity patterns within spikes imaged over ∼20 minutes. Together, our results show a considerable variation in dendritic spiking of L5 PNs, which strongly supports of the existence of several computational units within a dendritic tuft.

Our results show that branch variation coexists with global prevalence suggesting that prevalence is not a good measure of spike variation and, therefore, not a good measure for dendritic compartmentalization. Yet, most other studies that monitored activity in a substantial number of branches found a higher correlation in branch activity.

In agreement with recent studies, we found a low frequency of local dendritic activity ^6-8,20^. However, compared to these reports, we observed a higher degree of spiking variability. Technical features such as the number of branches imaged can explain this discrepancy. When a small number of branches per cell are imaged, they are typically closer to one another and of closer kin, therefore, displaying less variable activity ^16,20^. Underestimating variability could also arise from low expression levels of GECIs, which may miss weak events and thus over-represent broader and uniform spikes. High expression of GECIs may also increase the detection of widespread spikes, as it can increase the time course and the spatial spread of intracellular Ca^2+ 29,30^.

Several biological aspects affect the prevalence and variability of dendritic spikes. Some studies reporting a majority of global spikes focused on proximal apical dendrites (∼3^rd^ branching order) ^6,7^. At these locations, the strong influence of the soma, and the low signal attenuation caused by dendritic bifurcations, are likely to increase the incidence of strong, global, and coordinated spikes. Notably, the influence of the soma over dendritic activity can differ between cortical sites. For example, a recent study found that the somato-dendritic coupling of L5 PNs is higher at V1 than V2 ^31^. This finding is consistent with another study that found high somato-dendritic coupling in L5 PNs of V1 and that dendritic activity across branches was widespread and highly coordinated ^8^.

Dendritic activity is also affected by behavioral states (e.g., anesthetized, awake, and behaving). Experiments using isoflurane found that dendritic spikes in L5 PNs were broader under anesthesia than during wakefulness ^5^ or were uniform across multiple dendrites ^6^. Ketamine and xylazine also enhanced spiking ^32^. In the experiments presented here, mice had to balance their hindlimbs on a platform positioned above a moving rubber belt while striding their forelimbs over it. It is possible that during the acquisition of such a novel motor skill, dendrites receive more stochastic input that supports more variable dendritic spiking. It would be interesting to test whether the variability of dendritic activity changes as a function of practice.

Our finding that global spikes displayed differential branch activity is important for several reasons. First, the predominance of global spikes can be interpreted as an indication that apical tufts act primarily as a single computational unit. The abundance of glocal spikes suggests that global dendritic activity can coexist with local computations and supports the existence of multiple dendritic functional units. Second, glocal spikes are relevant for how local integration, which was thought to elicit mainly weak spikes, can affect the output of the remote soma. Glocal spikes offer a simple solution where spikes can maintain a mixture of global and local properties. Glocal spikes, therefore, undermine a strict distinction between global and local spikes. Although weak local spikes are limited in their ability to affect somatic output, these spikes can induce plasticity ^13,16,33^. Therefore, compartmentalized dendritic activity may lead to the differentiation of computational units across the tuft. In turn, global spikes could restrain such differentiation. It would be interesting to determine whether spikes of different prevalence have different effects on synaptic plasticity.

Glocal spikes are in line with two recent theoretical studies. The first found that the somatic output of L2/3 PNs could be predicted by the activity of linear dendritic subunits that elicit a global spike ^34^. The second concluded that tufts of L5 PNs contain several functional compartments, which are substantially fewer than dendritic branches ^35^. A direct assessment of the number of functional dendritic units would benefit from imaging tools that would enable imaging of an even greater number of branches.

Finally, it is essential to note that variable branch activity is not a property of apical dendrites alone. Studies of hippocampal PNs during navigation revealed that spiking activity of basal dendrites is highly variable and that dendrites can display place tuning independent of their soma ^36^. In addition, variable spiking activity was also observed in oblique dendrites of L5 PNs ^37^. Together, variable and dynamic dendritic activity is employed in all dendritic compartments, in many brain sites, and across a variety of behavioral states.

## Supporting information

Geron 2022 Video S1

Geron 2022 Video S2

Geron 2022 Video S3

## Acknowledgements

I thank Drs. Bernardo Rudy and Wenbiao Gan and their lab members for fruitful discussions; Drs. Orna Issler, Robert Machold, and Joseph Cichon for comments on the manuscript; Drs. Janani Sundararajan, Jason Moore, and Benjamin Schuman for help with MATLAB and statistical analysis; and Dr. Wei Li for assistance with experiments.

## Author contributions

With the help of the acknowledged individuals, EG designed research, conducted the experiments, analyzed the data, and wrote the manuscript.

## Materials and Methods

### Animals

Experiments involving the use of animals were in accordance with NIH guidelines and approved by the Institutional Animal Care & Use Committee. Male and female C57b mice at postnatal days (P) 30 were used for experiments. Only animals weighing more than 12 g were used. Animals were group-housed on a 12-hour light/dark cycle with light-on at 8 AM. Animals had free access to food and water. All experiments were initiated during the light-on period. *Cx3cr1*::eGFP mice were purchased from Jackson Laboratory (005582) and kept on a C57b background. WT C57b mice were purchased from Harlan Sprague Dawley Inc.

### Surgical procedures

#### Intracranial AAV injections of neonates

We used these viruses: AAV9.CaMKII::Cre, AAV9.FLEX.CAG.tdTomato, AAV9.FLEX.CAG.GCaMP6s (Addgene).

To maintain high titers of the injected viruses, we performed the following serial dilutions. The Cre-harboring virus was first diluted in ACSF (typically 1:700 to 1:1200, initial titer: 10^13^ VG/mL). This dilution was then diluted in ACSF that contained a fast-green dye (Sigma Aldrich, F725; to reach a final dye concentration of ∼ 0.1%, we first diluted a dye stock solution (6 % w/v) by 1:100, and then performed the subsequent dilutions). The Cre/dye mix was then diluted in the tdTomato-harboring virus at a ratio of 1:7 (7-fold excess of the tdTomato-harboring virus). Finally, the Cre/dye//tdTomato cocktail was mixed with the GCaMP-harboring virus at a 1:4 ratio (4-fold excess of the GCaMP virus). Therefore, the injected virus mix contained 80 % GCaMP-harboring virus, 17.5 % tdTomato-harboring virus, and ∼2.5 % ACSF (containing the Cre-harboring virus and the fast-green dye). A fresh virus mix was prepared for each day used.

For injections, a glass micropipette (Drummond 50001001×10) was pulled and beveled. A plunger was oiled lightly, inserted into the micropipette to pull the virus mix. Subsequently, pups at postnatal day 1-3 were anesthetized by hypothermia (typically ∼ 3 min on ice) and the micropipette was used (freehand) to penetrate the skin and skull and deliver 50-150 nL of the virus mix. M1 injection site was determined using the head’s veins as reference points (**Figure S1**).

#### Mounting a skull-attached head holder

∼24 hours before imaging, mice underwent surgery to attach a head holder. Specifically, mice were anesthetized with an intraperitoneal injection of ketamine and xylazine (100 and 10 mg/kg, respectively). The mouse head was shaved, and lidocaine (0.5 µg/µl) was applied to the shaved scalp. Subsequently, the skull surface was exposed, and the skull’s periosteum removed. Then, two metal bars were attached to the skull to the skull using a cyanoacrylate glue, and a well was molded from dental acrylic cement around the bars.

#### Installing a glass cranial window

After the head holder installation, a craniotomy was performed above the caudal forelimb in M1 ((AP: −0.5 ↔ +1 mm; DL: 1↔3 mm) ^38^. After removal of the skull, the exposed brain and meninges were immersed with artificial cerebrospinal fluid and covered with a glass coverslip. After completion, animals were placed on a heating pad until they regained mobility. We removed from the study animals that were hunched or scruffy, or lost 10% or more of their weight within a day after surgery.

### Behavior

#### Habituation to handling and head restraint

The mice were provided with wet food and habituated to handling for at least two weeks before surgery. Before imaging, the mice were given one day to recover from the surgery’s anesthesia and habituated to restraint in the imaging apparatus to minimize potential stress effects. Animals that resisted restraint (vocalizing or struggling for more than 20 s) were returned to their cage and tested again after more than 20 min.

#### Treadmill running

On the day of imaging, the animals were allowed to freely explore the imaging arena and treadmill for 5-10 min before restraining. Animals were then restrained and first imaged for structural data. The animals then receive an hour of rest and then imaged during acquisition of treadmill running. The treadmill speed was constant and ranged between 1.5-2.5 m per min (about 10 times slower than free running animals ^39^). A single running trial lasted for 2-4 minutes, interleaved by equal periods of rest.

### Two-photon imaging

#### Structural imaging

Imaging of neuronal structure was conducted using an Olympus Fluoview 1000 imaging system equipped with a 25× NA 1.05 water immersion lens. Neurons were imaged first at low magnification (1× zoom, 4 µm z step (∼1.5 µm pixel size)). Then higher magnifications were conducted to encompass the apical tuft (3× zoom, 2 µm z step (pixel size ∼0.5 µm)). Laser was supplied by a Ti:Sapphire laser (MaiTai DeepSee, Spectra Physics). Laser wavelength was tuned to 920 nm.

#### Volumetric imaging

Volumetric imaging was conducted on a custom-built microscope equipped with a Nikon 16× NA 0.8 water immersion lens ^26^. Imaging fields encompassed an area of ∼ 1 m^2^, where pixel size was 2.1 µm (typically 512 × 512 pixels). For each pixel, a depth of 40 µm was sampled quasi-simultaneously through 80 planes. Voxel dwell was 7.6 µs, and frame rate was 2 Hz. Laser was supplied by a Ti:Sapphire laser (MaiTai DeepSee, Spectra Physics). Laser wavelength was tuned to 935 nm. Laser power was adjusted to not exceed 30 mW/Cm^2^ at the sample. We found that continuous imaging exceeding 45 minutes did not induce apparent structural aberrations (not shown). We collected only the GCaMP channel during running.

### Image analysis

#### Ca^2+^ data extraction

Dendritic Ca^2+^ activity was measured by fluorescence changes of the indicators GCaMP6s. A maximal projection of all 80 z planes was created for each time point for image analysis of volumetric data. The resultant projections were then registered using the TurboReg plugin (Fiji) ^40^. Subsequently, regions-of-interest (ROIs) were created manually over an averaged projection of the registered time series to encompass the entire dendritic length (in branches longer than 25 μm). ROIs excluded intersections with other branches and passing axons (**Figures S3 A-D**).

We used a two-step process to determine F_0_. First, an estimated F_0_ was calculated as the 12^th^ percentile of the raw signal. This estimated F_0_ was used to compute an initial ΔF/F_0_. We then thresholded the original raw data to exclude measurements that yielded ΔF/F_0_ values below and above −10 and 20% (−10% to control for shifts in the z plane, and 20% to control for traces with high activity rates). The final F_0_ was the 12^th^ percentile of the thresholded raw data.

Spikes were defined as fluorescence elevation of 3 SDs above the baseline of the ΔF/F trace. F_0_ and the SD of the ΔF/F trace were determined individually for each branch. Spikes occurring within a time window of 1 s from other spikes were considered overlapping and were combined. For display of traces in figures 2-4, ΔF/F values below −0.05 % were zeroed.

#### Determining branch activity during spiking

The area under the curve (AUC) of the ΔF/F plot was calculated for each branch as long as one of the branches displayed activity above the 3 SDs threshold (equal period for all branches per spike). This yielded measurements for 14.4 ± 2.04 branches per neuron for every spike (35.3 ± 6.5 spikes per neuron; 9 neurons from 8 animals).

#### Calculating coefficient of variation between branches during a single spike

The coefficient of variation (CV) was derived by calculating the relative size of the standard deviation from the mean (SD/mean of the ΔF/F signal).

#### Cluster analysis

Ca^2+^ activity was averaged for the entire duration of the spike (equal number of branches for all spikes; for each spike-equal duration for all braches). Spikes were then clustered according to their branch activity (Ward’s method; Agglomerative hierarchical cluster tree (MATLAB)).

#### Evaluating the number of distinct activity patterns

We used four cluster evaluation indices: Davies Bouldin, Calinski Harabasz, Silhouette, and Gap (Figures 4C, D). For all indices, the linkage of the agglomerative clustering algorithm was determined by Ward’s method. Range of clusters was set to 1:6 (evalclusters; MATLAB).

### Statistical analysis

We used t-test to compare two groups and one-way ANOVA to compare more than two groups. One-way ANOVA tests were followed by Fisher’s LSD test for multiple comparisons. All tests were conducted as two-sided tests. Data are presented as mean ± SEM. Data is available upon request.

## Supplemental information

**Figure S1:**
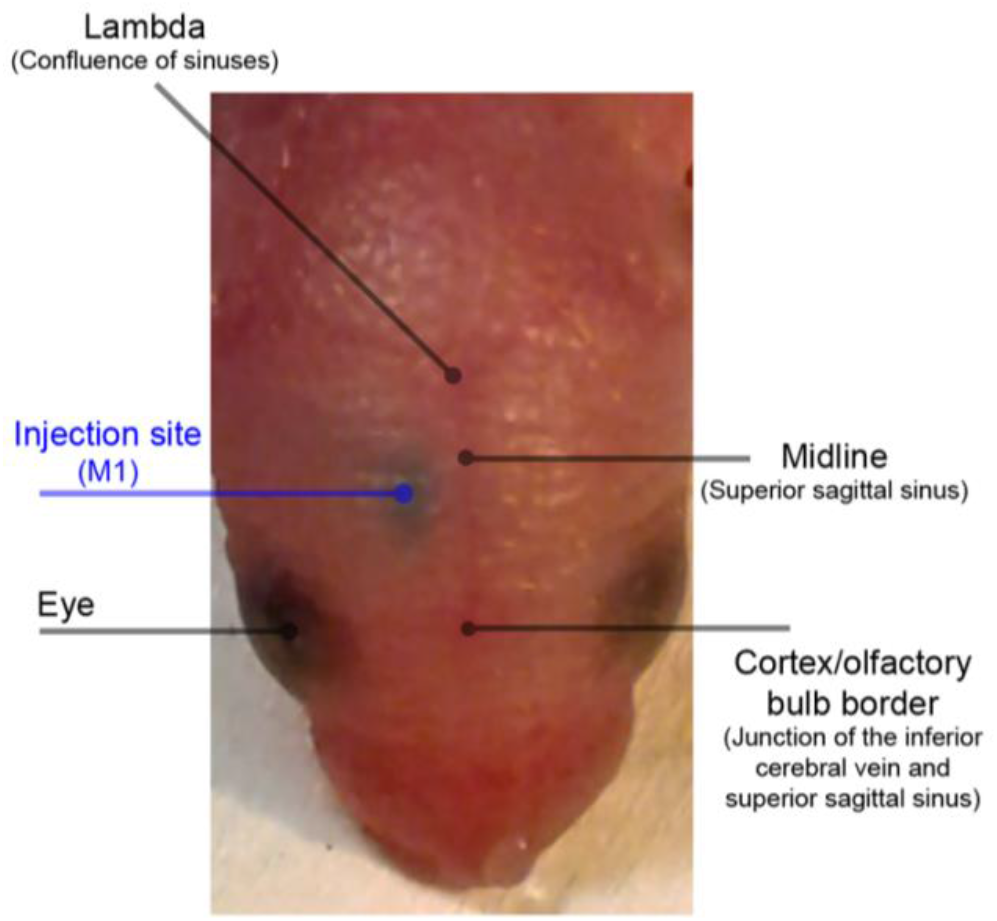
Using the head’s veins to target injection to the primary motor cortex. The neonate (P1) was anesthetized by hypothermia on ice, and the head’s veins served as reference points to target the primary motor cortex. The dye in the injected mix indicates the spread of the viral mix, and that the cortex was targeted correctly (as opposed to subcortical regions).

**Figure S2:**
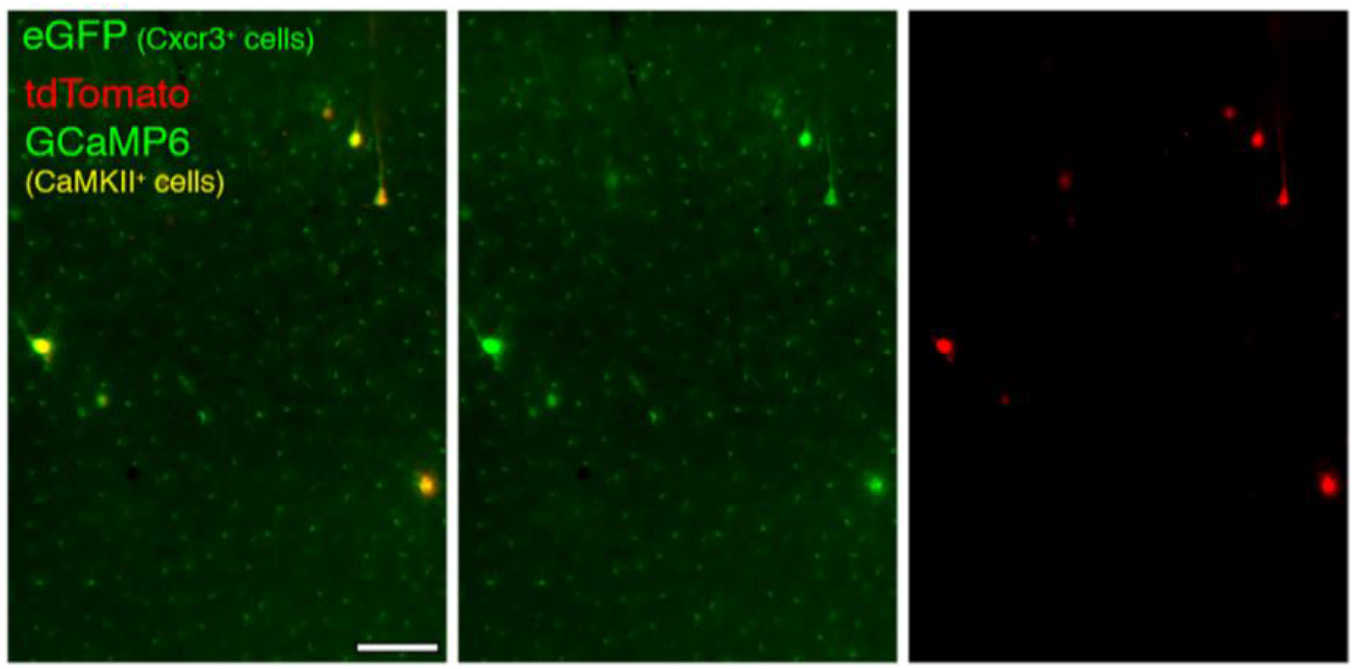
Stereotypical tiling of microglia in animals infected as pups. Microglia did not accumulate and were tiled evenly in animals infected with AAVs as pups. *Cx3cr1*::eGFP mice were injected as in Figure 1, and imaged at P30. Image is a representative brain slice example of 5 animals. Scale bar represents 100 μm.

**Figure S3:**
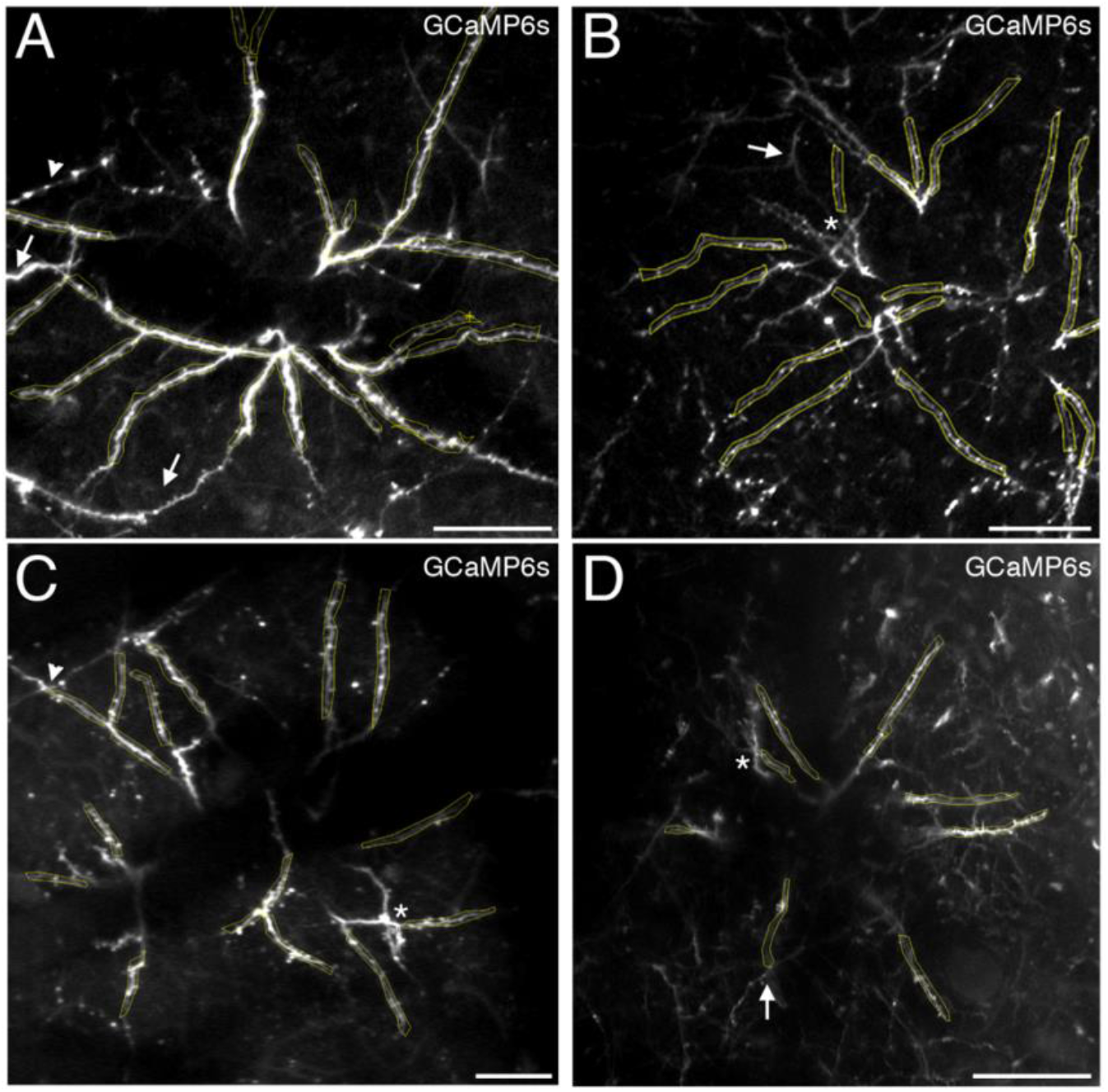
Examples of ROIs used in this study. A-D. Representative examples of ROIs used for dendritic Ca^2+^ data extraction. ROIs were created over an averaged projection of the registered time series to encompass the entire dendritic length. Markers over the panels denote the main guidelines used for ROI creation: exclude dendrites from neighboring cells (arrows), passing axons (arrowheads in A and C), and intersecting dendrites (asterisks). Images in A, B, and C correspond with the neurons presented in Figures 2, 3, and 4, respectively. Scale bras represent 50 μm in A and B, 40 μm in C, and 100 μm in D.

Video 1. **Dendritic Ca**^**2+**^ **spikes imaged in individual L5 PNs using volumetric imaging (example 1)**

This video shows 160 s of continuous Ca^2+^ imaging of dendrites of a GCaMP6s-expressing L5 PN during treadmill running. Time series here and in videos 2 and 3 are maximal projections of 80 z planes collected at 2 Hz. This video corresponds with the time projections and traces in Figure 2A, B. The video includes captions that point out Spike 1 and 2 presented in Figure 2B. It also includes captions that point out spikes of neighboring cells. Not all detected spikes were indicated.

Video 2. **Dendritic Ca**^**2+**^ **spikes imaged in individual L5 PNs using volumetric imaging (example 2)**

This video shows 64 s of continuous Ca^2+^ imaging of dendrites of a GCaMP6s-expressing L5 PN during treadmill running. This video corresponds with the time projections and traces in Figure 3A. The video includes captions that point out Spike 1 and 2 presented in Figure 3A. Not all detected spikes were indicated.

Video 3. **Dendritic Ca**^**2+**^ **spikes imaged in individual L5 PNs using volumetric imaging (example 3)**

This video shows 169 s of continuous Ca^2+^ imaging of dendrites of a GCaMP6s-expressing L5 PN during treadmill running. This video corresponds with the time projections and traces in Figure 4A. The video includes captions that point out Spike 1-3, presented in Figure 4A. Not all detected spikes were indicated.

**Table S1.** Structural data and spiking properties of the neurons imaged using volumetric microscopy. The data presented correspond to Figures 2-4. Neurons numbered 3, 6, and 7 are presented in Figures 2, 3, and 4 respectively.

|  | Depth of soma (μm under the pia) | Depth of main bifurcation (μm under the pia) | Number of branches analyzed | Bifurcation range (and median) | Distance (μm) and number of bifurcations between farthest imaged dendrites | Mean spike strength across all branches (% ΔF/F S<sup>-1</sup>; ± SEM) | Mean prevalence (± SEM) | Mean CV score (± SEM) | Number of spikes; Imaging time (min); Spiking rate |
| --- | --- | --- | --- | --- | --- | --- | --- | --- | --- |
| **1** | 625 | 135 | 9 | 3-5 (4) | 377, 8 | 73.9 ± 15.02 | 79.2 ± 4.4 | 65.1 ± 5.6 | 32; 13.2; 2.4 |
| **2** | 678 | 273 | 21 | 3-5 (5) | 570, 9 | 58.4 ± 7.5 | 82.1 ± 3.3 | 37.6 ± 4.7 | 43; 22.5; 1.9 |
| **3** | 670 | 270 | 27 | 3-6 (5) | 635, 10 | 61.4 ± 9.9 | 67.8 ± 3.3 | 41.9 ± 3.2 | 84; 25.5; 3.2 |
| **4** | 600 | 285 | 9 | 4-5 (5) | 527, 8 | 47.5 ± 5.1 | 75.4 ± 4.1 | 41.3 ± 2.1 | 46; 21.2; 2.1 |
| **5** | 680 | 390 | 11 | 3-6 (5) | 840, 9 | 24.1 ± 2.1 | 53.1 ± 6.5 | 50.07 ± 5.1 | 25; 21.4; 1.1 |
| **6** | 693 | 540 | 18 | 3-7 (5) | 1030, 11 | 32.8 ± 5.5 | 57.6 ± 7.2 | 54.7 ± 5.1 | 27; 21.4; 1.2 |
| **7** | 705 | 245 | 16 | 3-5 (5) | 486, 9 | 97.4 ± 16.1 | 67.5 ± 6.1 | 44.1 ± 1.1 | 25; 21.9; 1.1 |
| **8** | 620 | 210 | 8 | 4-5 (4.5) | 426, 9 | 54.9 ± 9.5 | 68.4 ± 7.3 | 40.2 ± 4.3 | 19; 13.5; 1.4 |
| **9** | 700 | 420 | 11 | 3-5 (4) | 805, 9 | 24.2 ± 3.03 | 51.4 ± 8.6 | 64.7 ± 9.6 | 17; 17.5; 0.96 |
| **Mean ± SEM** | 663.4 ± 12.8 | 307.5 ± 40.7 | 14.4 ± 2.04 | 3-7 (5) | 633.1 ± 72.1<br>9.1 ± 0.33 | 52.5 ± 7.5 | 67.1 ± 3.4 | 48.8 ± 3.3 | 35.3 ± 6.5;<br>19.8 ± 1.3;<br>1.7 ± 0.2 |

